# DORSSAA: Drug-target interactOmics Resource based on Stability/Solubility Alteration Assay

**DOI:** 10.1101/2023.12.29.573639

**Authors:** Ehsan Zangene, Elham Gholizadeh, Veit Schwämmle, Amir Ata Saei, Mathias Wilhelm, Lukas Käll, Mohieddin Jafari

**Author notes:** Corresponding author: Mohieddin Jafari.

## Abstract

Advancements in high-throughput techniques such as Thermal Proteome Profiling (TPP) and the high-throughput Proteome Integral Solubility Alteration (PISA) assay have revolutionized our understanding of drug-protein interactions. Despite these innovations, the absence of an integrative platform for cross-study analysis of stability and solubility alteration data represents a significant bottleneck. To address this gap, we introduce DORSSAA (Drug-target interactOmics Resource based on Stability/Solubility Alteration Assay), an interactive and expandable web-based platform for the systematic analysis and visualization of proteome stability and solubility alteration assay datasets. Currently, DORSSAA features 1,135,985 records spanning 38 cell lines and organisms, 135 compounds, and 480,456 potential protein targets. Through its user-friendly interface, the resource supports comparative drug-protein interaction analysis and facilitates the discovery of actionable therapeutic targets. Through two case studies; methotrexate target profiling in A549 cells and combinatorial-therapy drug–target interactions in leukemia cell lines, we demonstrate DORSSAA’s utility for identifying protein–drug interactions across diverse experimental contexts. This resource empowers researchers to accelerate drug discovery and enhance our understanding of protein behavior. Unlike data repositories and interaction knowledgebases, DORSSAA provides assay-native, protein-level MoA evidence with rigorous per-study FDR control, enabling context-specific target nomination, off-target deconvolution, and combination-aware discovery.

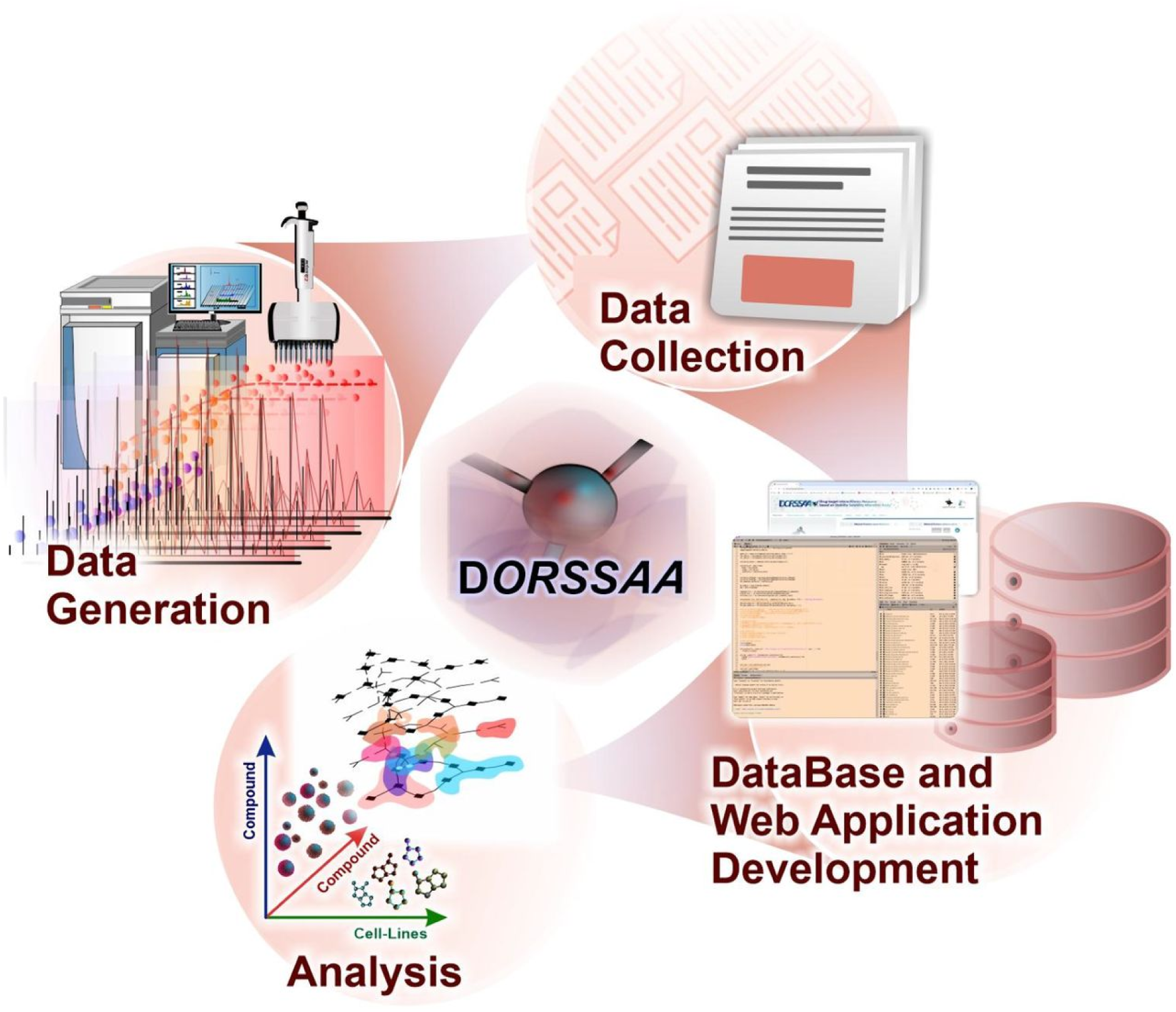

## Introduction

Drug discovery is a multidisciplinary field encompassing medicine, biotechnology, and pharmacology, aimed at identifying novel therapeutic compounds. Despite significant advances, a major challenge remains in the precise characterization of drug targets and off-targets within complex cellular environments. This challenge is especially pronounced in oncology, where identifying cancer vulnerabilities and dependencies, such as synthetic lethality relationships, is vital for developing precision therapies ^1,2,3,4^. This limitation is compounded by the absence of direct, systematic methods to monitor the effects of drugs on proteins within cells and tissues ^5^. Traditional evaluations by regulatory bodies such as the FDA, including absorption, distribution, metabolism, and excretion (ADME) studies and animal toxicity assessments, often fall short of fully elucidating a drug’s mechanism of action (MoA) ^6^. From a drug MoA point of view, identifying effective drug targets is challenging in drug development. As a result, many drug candidates fail during late-stage clinical trials (phase II trials), leading to resource wastage and underscoring the urgent need for innovative strategies to enhance drug development ^7^.

While small molecule modulators hold promise as therapeutic agents, their diverse effects on protein targets necessitate comprehensive profiling to better understand their molecular MoA. Although transcriptomic approaches like RNA sequencing and projects such as the Connectivity Map (CMAP) have been instrumental in mapping drug action ^8^. However, given that most FDA-approved drugs target proteins and the weak correlation between protein and RNA levels, protein-level changes are likely the most relevant readout of compound action ^9–12^.

Recent innovations in solubility and stability alteration assays have revolutionized the study of drug-protein interactions ^13,14^. These techniques, including cellular thermal shift assay (CETSA), thermal proteome profiling (TPP), and proteome integral solubility alteration (PISA) assay, leverage changes in protein stability or solubility upon ligand binding or environmental perturbations, offering an unbiased, high-throughput approach to probe drug interactions at the proteome level. By subjecting samples to varying conditions, these methods uncover critical insights into protein behavior and drug interactions across diverse biological contexts ^15,16^. Due to its numerous advantages, this approach has become widely used in both drug discovery and chemical proteomic research.

Centralized, queryable proteomics repositories are increasingly critical for nominating novel drug targets and dependencies, and for designing next-generation therapies. By enabling cross-dataset replication and mechanism-anchored readouts at the protein level, such resources help prioritize actionable targets, reveal context-specific vulnerabilities, and move beyond transcript-only inferences. Here, we present DORSSAA (Drug-target interactOmics Resource based on Stability Solubility Alteration Assay; https://dorssaa.it.helsinki.fi/), a comprehensive resource for the systematic analysis and visualization of solubility and stability alteration datasets. DORSSAA integrates high-throughput experimental data encompassing 1,135,985 records from 38 cell lines and organisms, 135 compounds, and 40,742 protein targets, providing an unprecedented dataset for drug interactomics. By consolidating this data into an intuitive, user-friendly web application, DORSSAA enables researchers to explore drug-target interactions across diverse experimental conditions. This resource not only facilitates cross-validation of findings but also accelerates the identification of actionable drug targets, enhancing the potential for translational drug discovery.

General proteomics repositories (e.g., PRIDE/ProteomeXchange) host raw and processed MS-based proteomics but do not harmonize thermal shift/solubility readouts into queryable, per-protein effect sizes across compounds and cell lines^17^. Drug-target knowledgebases (e.g., DrugBank, ChEMBL, BindingDB) curate biochemical/affinity interactions, rather than proteome-wide stability/solubility shifts^18^. Perturbational resources (e.g., LINCS/Connectivity Map) profile transcriptomic responses (L1000) instead of protein stability/solubility^8^. Drug-combination portals (e.g., DrugComb) focus on sensitivity/synergy metrics and do not provide protein-level thermal/solubility shift evidence^19^. DORSSAA offers several key advantages that make it a powerful complement to data repositories and interaction knowledgebases. It enables assay-native, protein-level harmonization of TPP, PISA, i-PISA, and CoPISA outputs into a unified relational schema. Its statistical framework ensures rigor through empirical DMSO-null p-values for PISA, per-study familywise false discovery rate control via Storey q-values with BY fallback, and safe p-value handling. DORSSAA also supports rich context modeling, incorporating cell-line and organism metadata, concentration and replicate tracking, and providing volcano and coverage views. Furthermore, it includes similarity layers—such as chemotype and cell-line similarity—to help generalize hit findings. Finally, its combination-aware evidence modeling in CoPISA facilitates rational pair design beyond classical synergy scores.

### Experimental Procedures

#### Collection of Proteomics Data

We assembled a harmonized corpus of 24 publicly available thermal-shift proteomics datasets, covering TPP, PISA, and i-PISA, together with our in-house CoPISA dataset^20^. In total, the resource spans 135 small molecules profiled across 38 cancer cell lines or species. Each study was reshaped to a tidy long format and enriched with consistent metadata (PMID, compound, cell line, UniProt ID, gene symbol, replicate, concentration, effect size, and statistical fields). Compound names were linked to PubChem InChIKeys, and cell lines were annotated with Cellosaurus and organism information. We then generated a coverage map (cell line × compound) and a normalized interaction fact table that captures per-protein stabilization/destabilization signals and their significance, and materialized everything into an indexed SQL backend for fast, app-ready queries in DORSSAA.

We curated a representative corpus of thermal-shift proteomics studies across assays, models, and temperature regimens (Table 1), which DORSSAA harmonizes for cross-study exploration.

**Table 1.** Studies curated for DORSSAA with assay, model/preparation, and temperature regime.

| No. | Study title | Assay | Model & preparation; Temperature (°C) | Reference |
| --- | --- | --- | --- | --- |
| 1 | Shifting Beyond Classical Drug Synergy in Combinatorial Therapy through Solubility Alterations | CoPISA | SKM-1, NOMO-1, MOLM-13, MOLM-16 (intact & extract); 48–59 °C | <sup>21</sup> |
| 2 | The antimicrobial drug pyrimethamine inhibits STAT3 transcriptional activity by targeting the enzyme dihydrofolate reductase | PISA | U3A (intact); 43–57 °C | <sup>22</sup> |
| 3 | Thermal proteome profiling for unbiased identification of direct and indirect drug targets using multiplexed quantitative mass spectrometry | TPP | K562 (intact); 37–67 °C | <sup>23</sup> |
| 4 | Lysine-specific demethylase 1A restricts ex vivo propagation of human HSCs and is a target of UM171 | TPP | HL-60 (intact); 37–67 °C | <sup>24</sup> |
| 5 | Thermal Proteome Profiling Identifies Oxidative-Dependent Inhibition of the Transcription of Major Oncogenes as a New Therapeutic Mechanism for Select Anticancer Compounds | TPP | MCF7 (intact); 37–67 °C | <sup>25</sup> |
| 6 | Anticancer Effect of Deuterium Depleted Water - Redox Disbalance Leads to Oxidative Stress | TPP | A549 (intact); 37–67 °C | <sup>26</sup> |
| 7 | Proteome Integral Solubility Alteration: A High-Throughput Proteomics Assay for Target Deconvolution | PISA | A549 (intact & extract); 43, 44.7, 46.4, 48.1, 49.8, 51.5, 53.2, 54.9, 56.6, 58 °C | <sup>15</sup> |
| 8 | Ion-Based Proteome-Integrated Solubility Alteration Assays for Systemwide Profiling of Protein-Molecule Interactions | I-PISA | A549, K562 (intact & extract); 37, 42, 46, 50, 54, 58, 62, 67 °C | <sup>27</sup> |
| 9 | A Simplified Thermal Proteome Profiling Approach to Screen Protein Targets of a Ligand | STPP | 293T (extract); 53–56, 66, 69, 72 °C | <sup>28</sup> |
| 10 | Thermal Proteome Profiling Reveals Glutathione Peroxidase 4 as the Target of the Autophagy Inducer Conophylline | TPP | U-2 OS (extract); 40, 43, 46, 49, 52, 55, 58, 61, 64, 67 °C | <sup>29</sup> |
| 11 | Thermal proteome profiling identifies PIP4K2A and ZADH2 as off-targets of Polo-like kinase 1 inhibitor volasertib | TPP | Jurkat (intact); 37, 41, 44, 47, 50, 53, 56, 59, 63, 67 °C | <sup>30</sup> |
| 12 | Thermal proteome profiling monitors ligand interactions with cellular membrane proteins | TPP | K562 (ATP), Jurkat (pervanadate) (intact & extract); 37–67 °C | <sup>31</sup> |
| 13 | Proteome-wide solubility and thermal stability profiling reveals distinct regulatory roles for ATP | TPP | Jurkat E6.1 (extract); 37–66.3 °C | <sup>32</sup> |
| 14 | Cell surface thermal proteome profiling tracks perturbations and drug targets on the plasma membrane | TPP | K562 (intact); 37–67 °C | <sup>33</sup> |
| 15 | Identifying the Target of an Antiparasitic Compound in Toxoplasma Using Thermal Proteome Profiling | TPP | Toxoplasma (CDPK1G, CDPK1M) (intact); 37–67 °C | <sup>34</sup> |
| 16 | Thermal proteome profiling of breast cancer cells reveals proteasomal activation by CDK4/6 inhibitor palbociclib | TPP | MCF7 (intact); 37–65 °C | <sup>35</sup> |
| 17 | Precipitate-Supported Thermal Proteome Profiling Coupled with Deep Learning for Comprehensive Screening of Drug Target Proteins | PSTPP | HEK293T, K562 (extract); 44, 52, 53, 65 °C | <sup>36</sup> |
| 18 | Thermal proteome profiling allows quantitative assessment of interactions between tetrachloroethene reductive dehalogenase and trichloroethene | TPP | Sulfurospirillum multivorans (extract); 43–97 °C | <sup>37</sup> |
| 19 | Thermal Proteome Profiling in Zebrafish Reveals Effects of Napabucasin on Retinoic Acid Metabolism | TPP | Zebrafish embryo (extract); 34–64 °C | <sup>38</sup> |
| 20 | Tracking cancer drugs in living cells by thermal profiling of the proteome | TPP | K562 (intact & extract); 37–67 °C | <sup>13</sup> |
| 21 | Systematic analysis of chemical-protein interactions from zebrafish embryo by proteome-wide thermal shift assay, bridging the gap between molecular interactions and toxicity pathways | PISA | Zebrafish embryo (extract); 37–67 °C | <sup>39</sup> |
| 22 | Identification of Celecoxib-Targeted Proteins Using Label-Free Thermal Proteome Profiling on Rat Hippocampus | TPP | Rat (extract); 67 °C | <sup>14</sup> |
| 23 | Network integration of thermal proteome profiling with multi-omics data decodes PARP inhibition | 2D-TPP | UWB1.289 (intact); 42.1, 44.1, 46.2, 48.1, 50.4, 51.9, 54, 56.1, 58.2, 60.1, 62.4, 63.9 °C | <sup>40</sup> |
| 24 | Large-scale characterization of drug mechanism of action using proteome-wide thermal shift assays | PISA | K562 (intact & extract); 48–58 °C | <sup>41</sup> |

**Table 2.** Database tables in DORSSAA, record counts, and where they appear in the app.

| Table name | # Records | Description (content & data types) | Shown in app tabs |
| --- | --- | --- | --- |
| Cell line | 38 | Unique cell lines with identifiers (character), Cellosaurus IDs (character), organism (factor/chr) | *protein/Cell-Line/compound tabs* |
| Compounds | 135 | Unique compounds (character); PubChem InChIKey (character) | *protein/Cell-Line/compound tabs* |
| DTC Target | 480,456 | Drug-target-common records use here as an external validation | *protein/Cell-Line/compound tabs* |
| Interaction | 1,135,985 | Fact table of protein-compound effects: UniProt, gene symbol, cell line, PMID, log2FC, p/FDR, dose | *protein/Cell-Line/compound tabs* |
| Target | 40,742 | Unique protein targets: UniProt ID; gene symbol | *protein/Cell-Line/compound tabs* |
| Cell lines similarity | 55 | Showing how different cell lines are similar based on different expressed biomarkers | *Cell-line tab* |
| Compound similarity | 528 | Pairwise compound similarity from SMILES fingerprints (drug1, drug2; similarity; numeric/char) | *Compound tab* |
| Drug target interaction | 19,378 | Curated drug-target interactions; records use here as an external validation | *protein/Cell-Line/compound tabs* |
| **Meltome** | 1,858,170 | Meltome/thermal stability records (species/protein) for each protein of interest | *protein/Cell-Line/compound tabs* |
| **Micha** | 367,628 | Micha compound annotations (IDs; primary targets; character) | *protein/Cell-Line/compound tabs* |

#### Data Normalization and Handling Missing Values

To ensure cross-study consistency, we standardized column names (replacing non-word separators with underscores), reconciled identifiers (prioritizing UniProt IDs and mapping HGNC symbols to UniProt via biomaRt when needed), and split multi-gene or multi-UniProt entries into separate rows. Inferred UniProt IDs are explicitly marked with a cloud symbol (☁) for transparency^42^.

Reported –log10(p) values were converted back to raw p-values where applicable, extreme/zero values were safely clamped, and multiple testing was controlled per (PMID × compound) family using Storey’s qvalue with a robust π₀ estimate, automatically falling back to Benjamini–Yekutieli for small or problematic families. For the K562 PISA study, we built an empirical null from DMSO controls (light row-mean imputation only when ≤20% missing), scored two-sided effects with add-one smoothing, and applied per-drug Benjamini-Hochberg False Discovery Rate (FDR). Replicate structure is preserved (with a dedicated replicate column and replicate-correlation QC), and variability across studies is visualized in DORSSAA via volcano plots so users can judge effect sizes in their experimental context.

#### Protein Query Database Interface

The Protein Query tab provides a versatile platform for users to investigate compounds treated within specific cellular contexts or organisms, enabling the identification of individual drug-target interactions or comprehensive database exploration. Query outputs are presented in multiple formats, including bar charts, volcano plots, and detailed data tables containing interaction-specific information. Users can further refine these results based on the significance thresholds of reported p-values.

Additionally, the interface facilitates cross-referencing of protein identities with UniProt records and provides direct access to external databases, such as DTC ^43^, Micha, and Drug Target Interact ^44^, for extended information on drug-target interactions ^43,45^.

#### Cell Lines Query Database Interface

The Cell Line Query tab offers a powerful platform for investigating critical biological questions by enabling users to explore the scope and extent of drug-protein interactions across various cellular contexts. By integrating insights from the Protein Query tab, users can evaluate which cancer cell lines or organisms are most conducive to drug-protein interactions and which are less responsive. This functionality provides valuable insights into pan-cancer and cancer-specific interactions, supporting the development of personalized therapeutic strategies. To enhance user experience and analysis capabilities, the interface includes advanced tools and features, such as interaction filtering based on p-values, UniProt protein annotations, external checks with UniProt, and cross-references to external drug-target interaction databases. These resources ensure comprehensive exploration and interpretation of interaction data within relevant biological contexts.

Additionally, the interface incorporates a melting points distribution tool sourced from the Meltome database ^46^. This tool allows users to visualize the melting point diversity of queried proteins across different contexts. Using box plots separated by cell line and a combined box plot for all cell lines, researchers can gain critical insights into melting point variations, providing a valuable perspective when designing experiments.

#### Compound query database interface

The Compound Query tab addresses critical aspects of drug repurposing by enabling users to investigate how a single protein can be targeted by multiple drugs. This feature highlights drugs with potential therapeutic applications against a specific protein within a defined cellular or organismal context, thus fostering innovative approaches to drug repurposing and therapy design.

Similar to the Cell Line Query tab, this interface includes a robust suite of analytical tools, including filtering interactions by p-values, accessing UniProt protein descriptions, performing external checks with UniProt, and verifying drug-target interactions through external databases. Additionally, we provided the same functionality mentioned in the Cell Line Query tab for protein melting points, allowing users to access insights into melting point diversity across different cell lines and contexts.

#### Protein Set Enrichment Tab

To enhance functional insights, the platform includes built-in tools for gene set enrichment analysis (GSEA) and singular enrichment analysis (SEA) based on Gene Ontology (GO) annotation ^47^. These tools enable the investigation of potential shared functional characteristics among proteins associated with specific compounds. The analysis is rooted in the hypothesis that proteins targeted by a given chemical may share molecular functions, biological processes, or cellular compartments as defined by GO terms. The results of enrichment analyses are visualized through treemaps and network diagrams, offering intuitive representations of interrelated GO terms with similar semantic features.

This framework further supports the hypothesis that compounds targeting distinct protein groups may exhibit functional similarities, reflected in shared gene ontologies. By incorporating semantic similarity tools, users can quantitatively assess and visualize these relationships, providing valuable insights into the functional and mechanistic overlap among drug targets.

#### Database Implementation

The DORSSAA backend server was developed using the R programming language (v.4.5.1) in conjunction with the Shiny App framework to provide an interactive web-based user interface. SQL database functionality was integrated via the DBI R package, enabling efficient data management and querying. JavaScript code was incorporated on both the front-end and server sides to enhance functionality and interactivity. Statistical analyses and data processing were conducted entirely in R. DORSSAA was rigorously tested for compatibility and performance across multiple widely-used web browsers, including Google Chrome (preferred), Firefox, and Apple Safari. The system ensures seamless user interaction regardless of the chosen platform. The design and development workflow for DORSSAA is illustrated in **Figure 1**.

**Figure 1:**
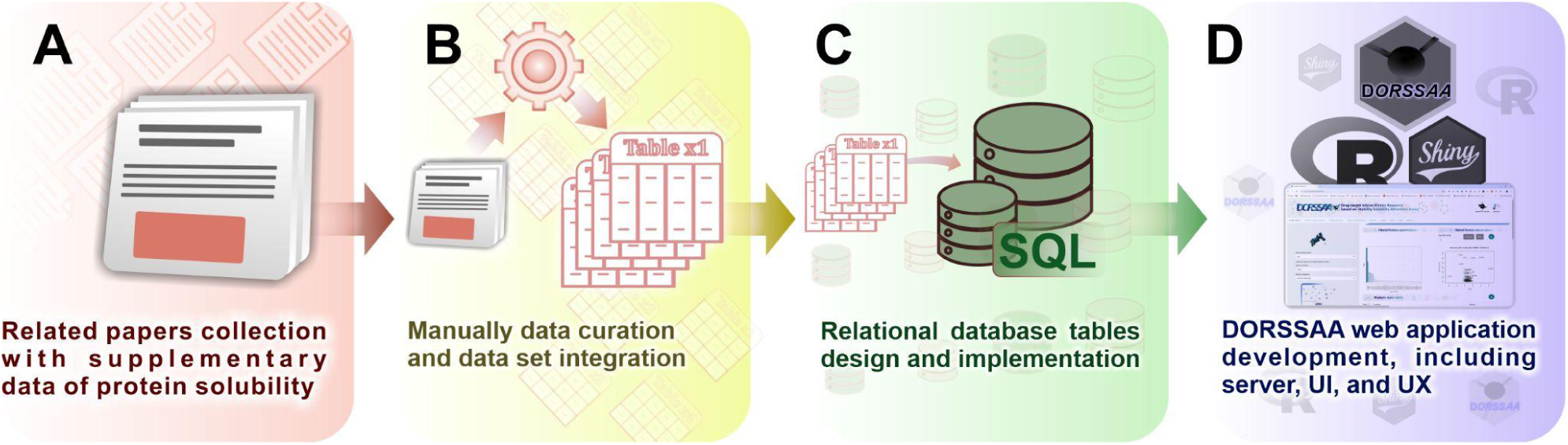
Schematic representation of the DORSSAA workflow. (A) Data collection from relevant publications. (B) Manual data curation and integration into datasets. (C) Relational database schema design. (D) Implementation and development of the DORSSAA web application.

#### Database Content

DORSSAA contains a comprehensive and diverse dataset tailored to the needs of researchers studying drug-protein interactions:

A. **Number of Records**: The core “Drug-Protein Interaction” table includes 1,135,985 entries, representing interactions derived from 38 distinct cell lines and organisms, 135 compounds, and 40,742 proteins (as of 25 October 2025).
B. **Data Accessibility**: Researchers can interact with the data through the DORSSAA web application, an intuitive and interactive web-based interface available at https://dorssaa.it.helsinki.fi/. The interface allows users to retrieve results for any drug-target interaction based on proteome solubility assay data.
C. **Ongoing Updates**: The database is updated regularly as new publications with publicly available TPP datasets are released. This ensures that DORSSAA remains a current and reliable resource for the scientific community.
D. **Long-term Availability**: The graphical user interface (GUI) offers free access to researchers and is committed to indefinite availability, making DORSSAA a dependable and enduring tool for drug discovery and development investigations.

#### Web interface

The DORSSAA web application offers an intuitive and interactive platform for data retrieval, visualization, and export (**Figure 2**). The interface is structured around three main functional tabs: Protein Query, Cell Line Query, and Compound Query backed by additional analysis and utility views including Protein Set Enrichment, Statistics, Studies, News, Help, and About us.. Users can initiate their queries by selecting options from three drop-down menus: UniProt_ID or Gene_symbol, Cell Line, and Compound. Upon selection, the main panel renders a bar chart and a volcano plot, and a detailed table summarizes all relevant interactions with comprehensive annotations and metrics (**Figure 2**). These visual and tabular outputs let users examine drug–target interactions, assess protein-level changes, and extract insights into molecular mechanisms.

**Figure 2:**
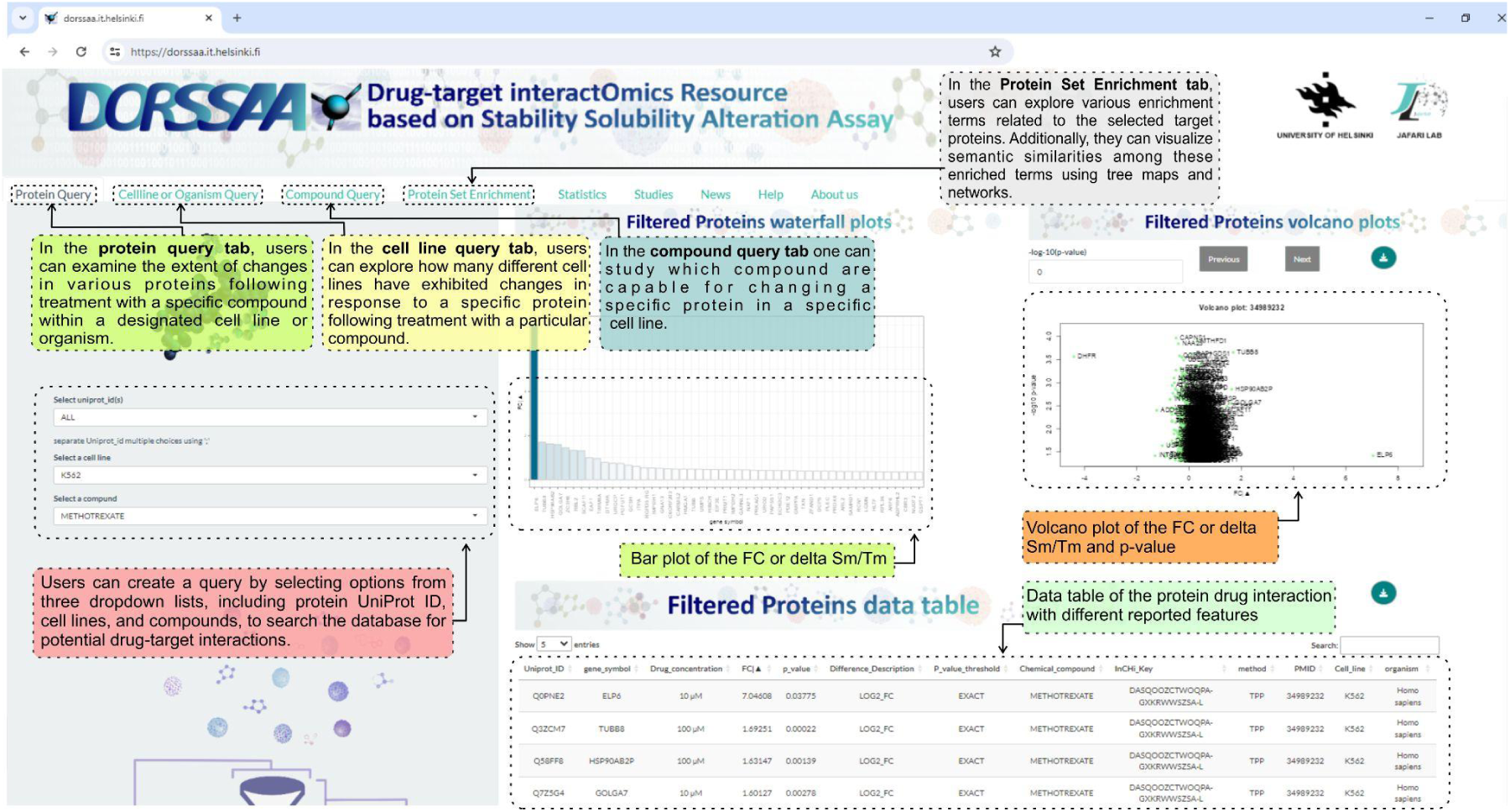
The DORSSAA web application showcases an intuitive interface divided into three primary tabs: Protein Query, Cell Line Query, and Compound Query. Users can initiate queries by selecting options from dropdown menus for UniProt_IDs, cell lines, and compounds. The main panel features visualizations such as bar charts and volcano plots, offering insights into drug-target interactions and protein-level changes. Below the visualizations, a detailed data table provides comprehensive information on all identified interactions, supporting efficient data exploration and in-depth analysis.

Importantly, DORSSAA has undergone regular versioned releases to expand data coverage and functionality and to improve usability. Notable milestones include the addition of Cell line/organism search (v2.1.2), Compound Query and GSEA (v2.2.1), Meltome integration (v2.2.2), zebrafish and rat datasets (v2.2.3), FDR filtering and new datasets (v4.1.1), and a new database-search GUI in the Statistics page with broader UI/Server upgrades plus updated Help/About (v4.1.2). Performance-focused updates (v3.1.1–3.2.1) optimized SQL/server layers and streamlined drop-downs for large queries. This active maintenance and roadmap reflect our commitment to making DORSSAA the primary resource for stability/solubility-based interactomics. *(See Release History Table for full version notes in the News tab.)*

## Results

### Case Study 1: Exploring Methotrexate’s Protein Targets and MoA in A-549 Cells Using DORSSAA

To evaluate the accuracy and relevance of DORSSAA’s data, we conducted an in-depth analysis focusing on methotrexate’s protein targets within the A-549 cell line. Applying stringent filtering criteria, including a FDR threshold of 0.05, we identified statistically significant and biologically relevant interactions.

Using DORSSAA’s Protein Query tool, we compiled a detailed list of proteins associated with methotrexate. Cross-referencing with UniProt and other external databases showed strong concordance, supporting the reliability of DORSSAA’s outputs. Gene Ontology enrichment then highlighted six distinct functional categories, with notable over-representation of oxidative-stress–related biological processes.

Among the 15 target records identified, Dihydrofolate Reductase (DHFR) (UniProt ID: P00374) emerged as a top target. DHFR exhibited the highest log2-fold change (log2-FC), with a highly significant of FDR, underscoring its strong association with methotrexate in A-549 cells (**Figure 3A**). DHFR is the cognate target of methotrexate. DHFR is known for its role in the biosynthesis of amino acids and folic acid, reducing dihydrofolate to tetrahydrofolate via the nicotinamide dinucleotide phosphate cofactor. Its therapeutic relevance in cancer and bacterial infections is well-documented. Interestingly, in DORSSAA, DHFR was targeted by five additional drugs (e.g., staurosporine, SNS032, raltitrexed, and panobinostat) across different studies, as shown in the Compound Query tab (**Figure 3B**).

**Figure 3.**
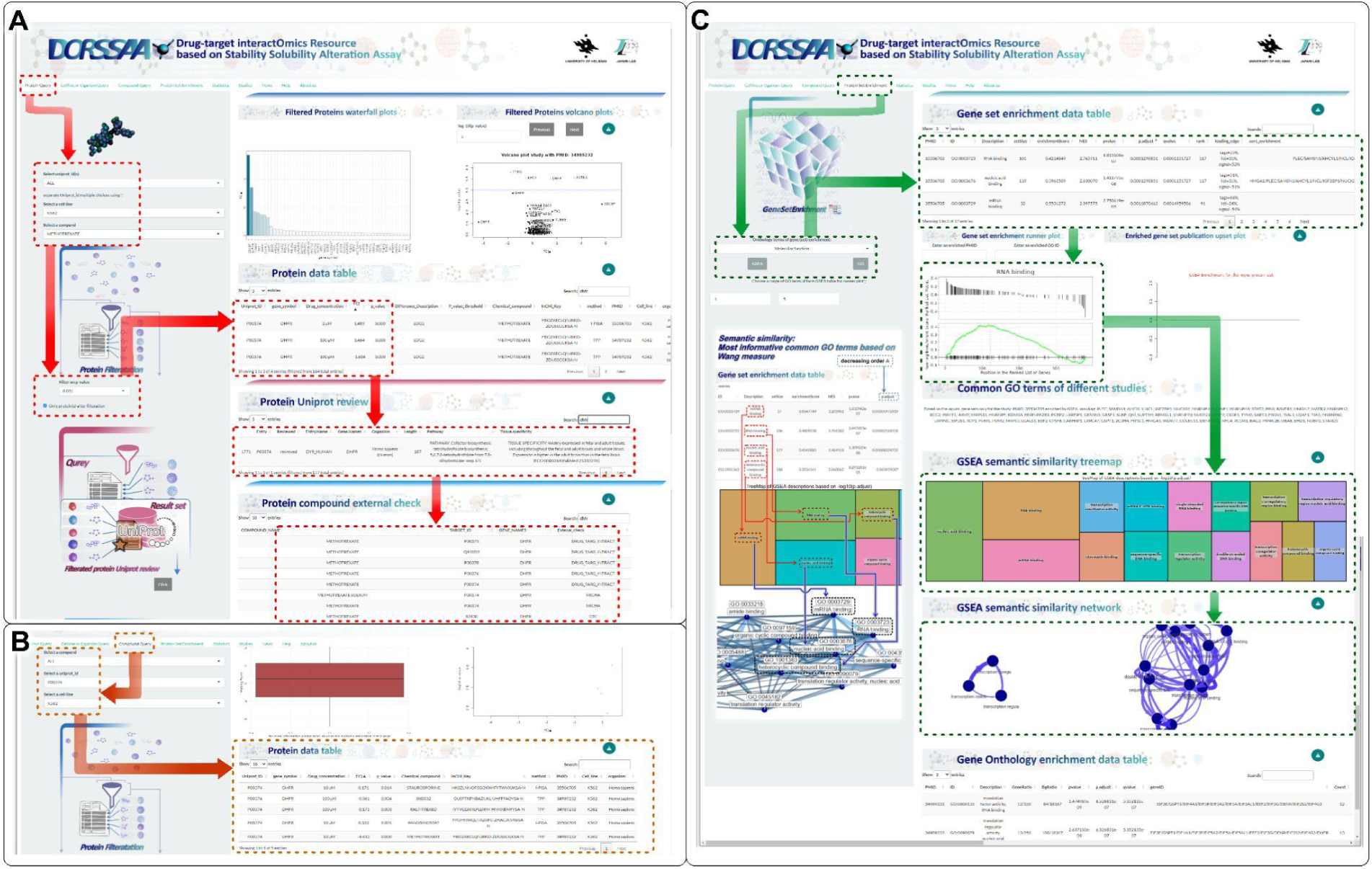
Methotrexate-Associated Proteins and DHFR Target Identification Using DORSSAA. (A) Protein Query analysis identified 13 methotrexate-associated proteins, with DHFR showing the highest log2-fold change and significance. (B) DHFR was also targeted by five additional drugs, as shown in the Compound Query tab. (C) Enrichment analysis and cross-referenced studies confirmed DHFR’s central role in methotrexate’s MoA and highlighted opportunities for drug repurposing and combinatorial therapy.

To explore methotrexate’s MoA, biological process enrichment analysis revealed significant terms, including its inhibition of transmethylation reactions. This interaction was further validated across multiple studies using techniques like TPP and Single-tube Thermal Proteome Profiling (STPP), spanning cell lines such as K562 and 293T. Notably, while K562 cells are derived from chronic myelogenous leukemia, 293T cells originate from embryonic kidney cells, suggesting a degree of universality in DHFR-methotrexate interactions. Furthermore, DORSSAA enabled the identification of drug repurposing opportunities for targeting DHFR using other compounds, highlighting its potential as a target for combinatorial and personalized therapeutic strategies (**Figure 3C**).

### Case Study 2: Drug-Protein Interactions in Molm16 Cells Investigated Using DORSSAA

A complete submatrix inside the DORSSAA dataset (6 compounds × 4 cell lines) is matched to our recent studies of combinatorial therapy on 4 different AML cell lines with 4 single drugs and 2 combinations ^48,21,49^. In this case study, we leveraged DORSSAA’s unique analytical capabilities to investigate context-specific drug-target interactions and their combinatorial effects.

To begin, we analyzed the effects of sapanisertib on the Molm16 cell line. In the Protein Query tab, significant targets were identified by applying a FDR threshold of 0.05. Among the resulting targets, H2AFX (UniProt ID: P16104) stood out with a log-fold change of -1.673 and a FDR of 0.03993, indicating significant drug-induced alterations compared to untreated controls (**Figure 4A**).

**Figure 4:**
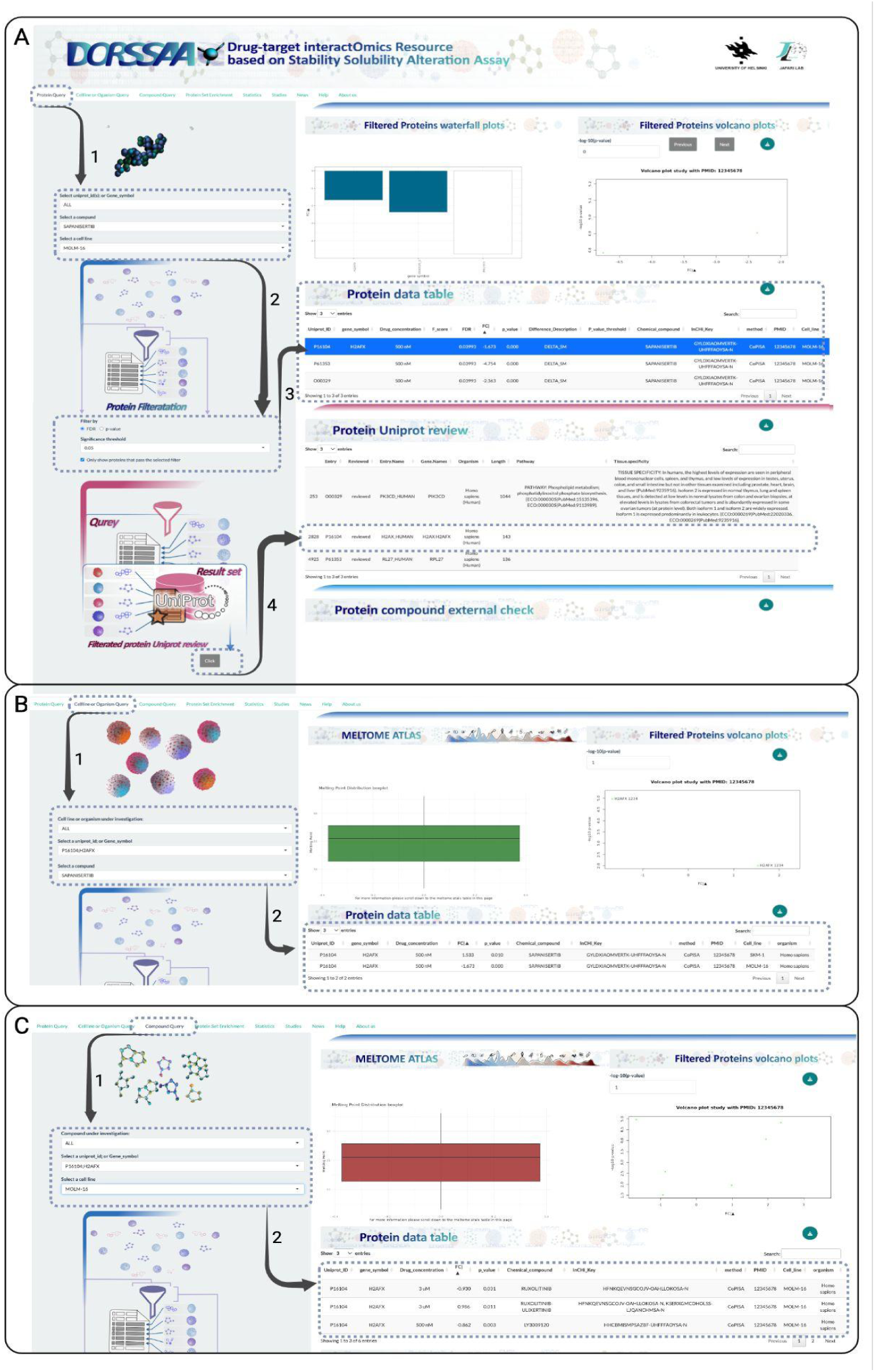
Drug-target Interactions in Molm16 Cells Using DORSSAA. (A) In MOLM-16 cells, sapanisertib significantly altered H2AFX (UniProt P16104; log₂ fold change = −1.673; FDR = 0.03993). (B) In SKM-1 cells, sapanisertib also targeted H2AFX (log₂ fold change = +1.533; p = 0.010). (C) In MOLM-16, H2AFX is additionally modulated by other single agents and combinations, including Ruxolitinib–Ulixertinib, LY3009120, Ruxolitinib, and Ulixertinib; each with distinct fold-change and significance value.

In the Cell Line/Organism Query tab, we assessed cell-type specificity across lines and cancer subtypes. The interaction was significant only in MOLM-16 and SKM-1, indicating an AML-context–specific relationship with potential implications for precision therapy (Figure 4B). In the Compound Query tab, additional agents targeting H2AFX emerged, Ruxolitinib–Ulixertinib, LY3009120, Ruxolitinib, and Ulixertinib, with varying log₂ fold changes and p-values. Notably, the Ruxolitinib + Ulixertinib combination showed an amplified effect on H2AFX (log₂ fold change = 0.986; p = 0.011) (Figure 4C). Collectively, DORSSAA’s integrated views enable researchers to traverse targets, cell contexts, and compounds, revealing novel relationships and actionable insights ^48,50^.

This comprehensive case study underscores DORSSAA’s strength in enabling the identification and exploration of both individual and combinatorial drug-target interactions, providing valuable guidance for targeted drug development and therapeutic innovation.

## Discussion and conclusion

DORSSAA serves as a comprehensive and purpose-built resource for drug discovery and protein research, enabling systematic assessment of cellular-level target engagement and off-target binding. By integrating advanced proteomics techniques, namely TPP and PISA, DORSSAA harnesses the power of mass spectrometry-based proteomics to characterize drug-protein interactions with high resolution and biological relevance.

Unlike transcript-only proxies, DORSSAA provides assay-aware, protein-level evidence to address critical questions in drug-target interactomics. These include: (i) Context-specific MoA mapping, identifying proteins stabilized or destabilized by compounds in defined cellular or organismal contexts; (ii) Off-target deconvolution, detecting non-canonical targets that shift consistently across studies; (iii) Target-centric querying, finding compounds that reproducibly modulate specific proteins or protein families across models; (iv) Cross-study replication, evaluating whether effect directions and magnitudes are consistent across independent assays and batches; (v) Combination design, distinguishing protein shifts unique to drug pairs versus single agents; (vi) Translational prioritization, identifying candidates that meet stringent FDR criteria, replicate across datasets, and map to tractable biology for follow-up; (vii) Generalizability vs. subtype specificity, determining whether target–compound interactions are broadly conserved or restricted to specific cancer subtypes; and (viii) Drug mimicry and repurposing, discovering compounds that recapitulate a target–compound’s thermal stability signature and may be rationally repurposed.

The core objective of DORSSAA is to accelerate therapeutic innovation and deepen our understanding of protein behavior under diverse conditions. Its extensive database compiles differential proteome solubility datasets from a wide range of sources, offering a rich and diverse proteomic landscape for exploration. The accompanying web application enhances accessibility through intuitive tools for visualizing, analyzing, and interpreting drug-target interactions, bridging the gap between complex datasets and actionable insights.

One of DORSSAA’s key strengths lies in its ability to support both focused and broad biological inquiries. Researchers can investigate how specific proteins respond to compound treatments across various cell lines and organisms, while also uncovering broader patterns of proteomic change under different experimental contexts. The compound query functionality further enables identification of drugs that may affect particular proteins, supporting drug repurposing and the design of combination therapies.

By providing orthogonal, assay-native signals ^51^, DORSSAA complements large-scale perturbation screens and refines cancer-dependency discovery ^1^. It also informs the design and prioritization of synthetic-lethal strategies ^2^, offering a valuable layer of evidence for translational research.

Technically, DORSSAA is built on a robust foundation using R programming, SQL database functions, and JavaScript, ensuring a seamless and reliable user experience across major web browsers. The database currently includes data from 37 distinct cell lines and organisms, 39 unique compounds, and 40,004 protein targets—making it a highly versatile and expandable resource.

To remain relevant in the fast-evolving landscape of proteomics and drug discovery, DORSSAA is committed to continuous updates. By promptly integrating newly published, publicly available datasets, it ensures that users have access to the most current and comprehensive information. This dedication to staying current reinforces DORSSAA’s role as an essential tool for researchers tackling complex challenges in drug discovery and protein science.

## Acknowledgment

We sincerely thank Bernhard Küster for his valuable feedback and insightful suggestions, which significantly contributed to improving both the manuscript and the development of DORSSAA. We also acknowledge Salma Ibourki for her assistance with data collection during her summer visit to our lab.

## Author Contributions

E.Z. contributed to the project, including data collection, design, and implementation of the SQL database, development of the server and UI/UX components, and oversight of the graphical aspects of the web application. E.Z. also managed deployment on the University of Helsinki’s VPS Linux server and led the manuscript writing. E.G. contributed to data collection and assisted in writing the manuscript. V.S., A.A.S., M.W., and L.K. provided valuable and constructive feedback on both the manuscript and the web application. M.J. conceived the research concept, provided supervision and coordination of the project, and offered constructive feedback on the manuscript and web application development.

## Funding

This study was financially supported by the Research Council of Finland [Grant 332454 to M.J.], Jane and Aatos Erkko Foundation [Grant 220031 to M.J.], Swedish Research Council [Grant 2020-00687 to A.A.S.], and the Swedish Society of Medicine [Grant SLS-961262, 1086 Stiftelsen Albert Nilssons forskningsfond to A.A.S.]. EZ’s salary was partially supported by the iCANPOD postdoctoral program, which is funded through the iCANDOC doctoral education pilot in precision cancer medicine..

## Conflict of interest

The authors declare no conflicts of interest.

## Notes

https://dorssaa.it.helsinki.fi/

### Competing Interest Statement

The authors have declared no competing interest.

### Summary of Updates

App: New DORSSAA release is live (https://dorssaa.it.helsinki.fi/): 1,135,985 interactions, 38 cell lines/organisms, 135 compounds, 40,742 proteins. Faster backend, new datasets, updated Statistics page. Manuscript: Updated author list, harmonized counts, clearer stats, more analysis, expanded study tables, tightened Protein/Cell/Compound Query sections, refreshed case studies (MTX→DHFR; AML H2AFX incl. Ruxo+Ulix), cleaner figures/captions, and references.

